# An ecological assessment of the pandemic threat of Zika virus

**DOI:** 10.1101/040386

**Authors:** Colin J. Carlson, Eric R. Dougherty, Wayne Getz

## Abstract

The current outbreak of Zika virus poses a threat of unknown magnitude to human health^1^. While the range of the virus has been cataloged growing slowly over the last 50 years, the recent explosive expansion in the Americas indicates that the full potential distribution of Zika remains uncertain^2-4^. Moreover, most current epidemiology relies on its similarities to dengue fever, a phylogenetically closely related disease of unknown similarity in spatial range or ecological niche^5,6^. Here we compile the first spatially explicit global occurrence dataset from Zika viral surveillance and serological surveys, and construct ecological niche models to test basic hypotheses about its spread and potential establishment. The hypothesis that the outbreak of cases in Mexico and North America are anomalous and outside the ecological niche of the disease, and may be linked to El Nino or similar climatic events, remains plausible at this time^7^. Comparison of the Zika niche against the known distribution of dengue fever suggests that Zika is more constrained by the seasonality of precipitation and diurnal temperature fluctuations, likely confining the disease to the tropics outside of pandemic scenarios. Projecting the range of the diseases in conjunction with vector species (*Aedes africanus*, *Ae. aegypti*, and *Ae. albopictus*) that transmit the pathogens, under climate change, suggests that Zika has potential for northward expansion; but, based on current knowledge, Zika is unlikely to fill the full range its vectors occupy. With recent sexual transmission of the virus known to have occurred in the United States, we caution that our results only apply to the vector-borne aspect of the disease, and while the threat of a mosquito-carried Zika pandemic may be overstated in the media, other transmission modes of the virus may emerge and facilitate naturalization worldwide.

## Main Text

Following a twenty-fold upsurge in microcephalic newborns in Brazil tentatively linked to Zika virus (ZIKV), the World Health Organization has declared an international health emergency^1^. Despite being profiled for the first time in 1947^8^, Zika remains poorly characterized at a global scale. Thus, the present pandemic expansion in the Americas poses a threat of currently unknown magnitude. Closely related to dengue fever, Zika conventionally presents as a mild infection, with 80% of cases estimated to be asymptomatic^9^. The cryptic nature of infection has resulted in sporadic documentation of the disease and rarely includes spatially explicit information beyond the regional scale^1-4^. This greatly limits the confidence with which statistical inferences can be made about the expansion of the virus. With an estimated 440,000-1,300,000 cases in Brazil in 2015^9^, and continuing emergence of new cases in Central America and, most recently, the United States, assessing the full pandemic potential of the virus is an urgent task with major ramifications for global health policy.

Current evidence portrays the global spread of ZIKV as a basic diffusion process facilitated by human and mosquito movement, a hypothesis supported by the frequency of infected traveler case studies in the Zika literature^10-13^. Tracing phylogenetic and epidemiological data has revealed the expansion of ZIKV has occurred in a stepwise process through the South Pacific, moving the disease from Southeast Asia into French Polynesia and the Philippines, and subsequently to Easter Island^1-4^. ZIKV is conjectured to have dispersed into South America as recently as three years ago from the last of those locations, and the virus is not presumed to be at a biogeographic equilibrium in the Americas. With cases in the ongoing outbreak in Colombia, El Salvador, Guatemala, Paraguay, and Venezuela, and by November of last year, as far north as Mexico, Puerto Rico, and the continental United States, the full potential distribution of the disease remains unknown. Moreover, alternative explanations for the disease’s expansion remain unconsidered; most notably, the role of climate change in Zika’s expansion is uncertain^7^.

We present three competing hypotheses that describe the path of expansion that Zika could take, based on evaluations of the ecological niche of the virus within and outside of its vectors. If the Zika niche is indistinguishable from that of its *Aedes* vectors (as is essentially the case for dengue fever^14^), future range expansions should match mosquito ranges. On the other hand, if Zika has a transmission niche that is constrained by climatic factors within the ranges of its mosquito vectors, its range may be much more limited—with, as we show below, possible confinement to the tropics. In this case, the expansion of Zika into North America represents one of two hypothesized processes: a steady range expansion driven by climatic shifts, or an anomalous event driven by human dispersal or extreme weather events. To test these hypotheses, we present the first spatially explicit database of Zika occurrences from the literature and an ecological niche model^15^ using that data to map the potential distribution of the virus.

Our dataset includes 64 of the known occurrences of the disease – a combination of clinical cases and seropositivity surveys in humans and mosquitoes. Of these, 60 points from outside the current outbreak are used in our model to determine the expected distribution in the Americas based on the niche in areas where the virus is established (rather than potentially transient). Spanning seven decades, these data have not previously been compiled nor explicitly geo-referenced, and emergency modeling efforts for diseases of special concern often work from limited data. Previously published sensitivity analyses unequivocally suggest that accuracy of the modeling methods we employ plateaus at or near 50 points, justifying the use of a dataset of this size^16-18^. Ensemble modeling also vastly improves the predictive power with datasets of this sort (Extended Data Fig. 1-5), and reduces the associated error. Our final model combines seven methods with a variable set chosen from bioclimatic variables and a vegetation index to minimize predictor covariance. The ensemble model performs very well (AUC = 0.993; Fig. 1), and strongly matches most occurrences including the hotspots of Brazilian microcephaly. It also predicts additional regions where Zika is as yet unrecorded, but where further inquiry may be desired (in particular, Southern Sudan and the northern coast of Australia). Our model indicates that certain occurrences, like the 1954 report from Egypt and almost all North American cases, are likely outside the stable transmission niche^19^ (i.e., persistent over time) of the virus. Moreover, we note that visual presentation of cases at the country level may make the range of the virus appear far larger than our models suggest (see Fig. 1). Projecting niche models to the year 2050 suggests that expansion of Zika’s niche outside the tropics is an unlikely scenario, independent of vector availability (Fig. 2d). However, significant westward expansion in South America and eastward expansion in Africa implies that Zika may continue to emerge in the tropics.

**Figure 1.**
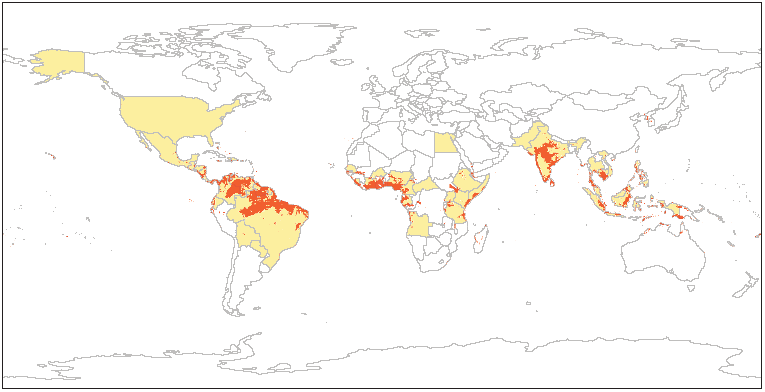
The global distribution of case reports of Zika virus (1947 to February 2016) broken down by country (blue shading) and an ensemble niche model built from occurrence data (red shading). Our model predicts occurrence in part of every shaded country; it is clear that displaying cases at country resolution overstates the distribution of the virus, especially in the Americas (for example, Alaska).

**Figure 2.**
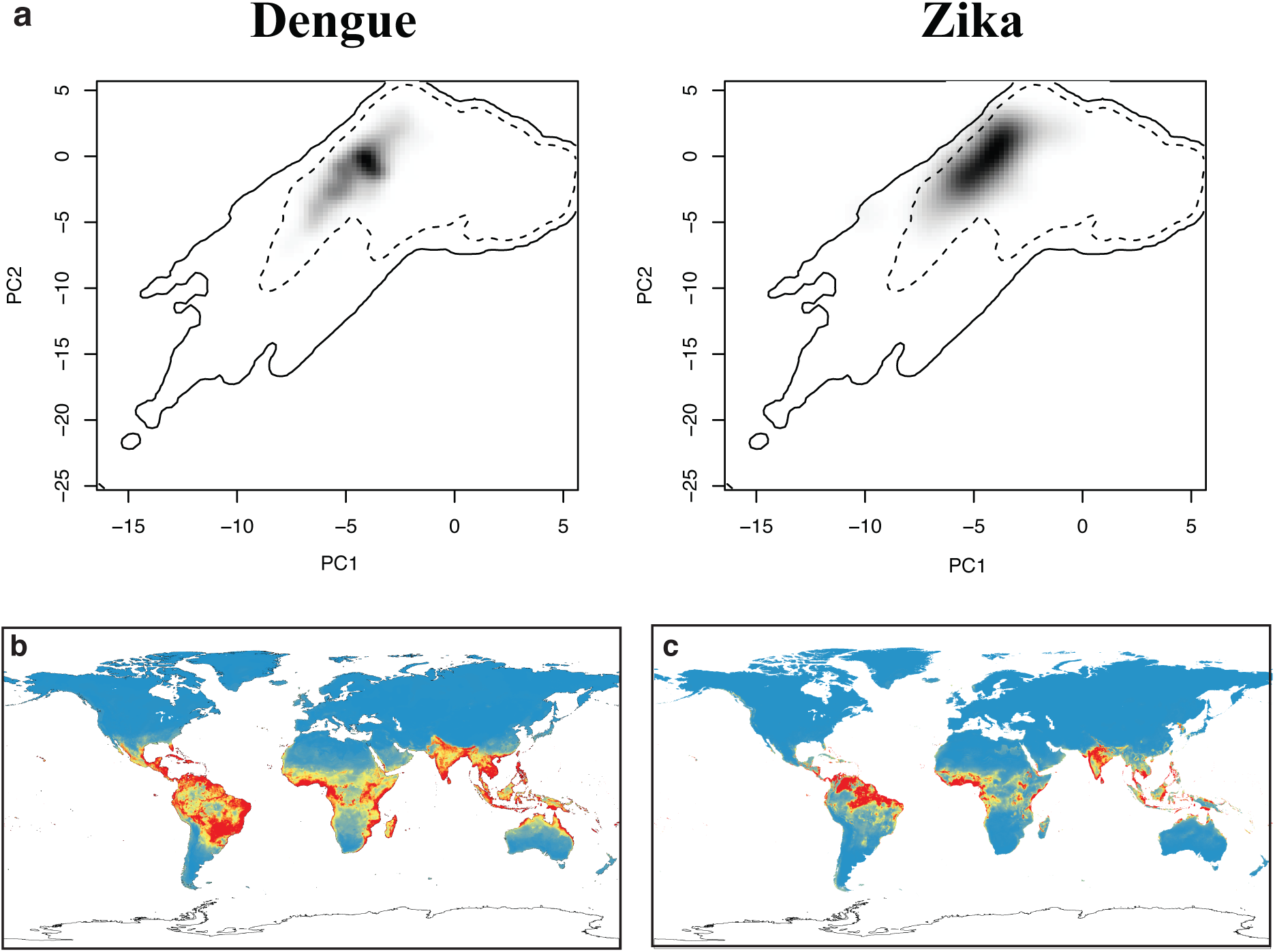
The ecological niche of Zika and dengue in principal component space (a). Solid and dashed lines are 100%/50% boundaries for all environmental data Despite apparent overlap in environmental niche space, the dissimilarity between the black shading in each principal component graph indicates statistically significant differences between the niches, evident in the projections of our niche models for dengue (b) and Zika (c).

Given the public health crisis posed by Zika, and the potential costs associated with underpredicting the extent of the current outbreak, we pay special attention to evaluating the sensitivity of our models to variations in our preliminary dataset. Geographical data on cases in the Americas are lacking, and the routes and drivers of transmission involved in that outbreak are uncertain, preventing cross validation of models of the current outbreak with our Old World model. But, in light evidence that African and Asian strains of the virus may be ecologically distinct, we present models trained on each continent and projected globally, as a basic sensitivity analysis (Extended Data Fig. 7). The two models cross-validate poorly; driven by both the 50% reduction in sample size and the higher degree of aggregation of Asian occurrences, the two projected distributions are severely different. But despite the major over-prediction of the Asian model and the overfitting of the African model, we emphasize that neither extreme scenario predicts any greater range in the Americas. Moreover, despite low transferability between continents, both sub-models are well matched by our aggregated model in their native range, further supporting the accuracy and predictive power of our global projection.

Recently published work by Bogoch *et al.*^6^ uses an ecological niche model for dengue as a proxy for the potential full distribution of ZIKV in the Americas, presenting findings in terms of potential seasonal vs. full-year transmission zones. While that approach has been effectively validated for dengue transmission in mosquitoes, using a model of one disease to represent the potential distribution of another emerging pathogen is only a placeholder, and is particularly concerning given the lack of evidence in our models that ZIKV and dengue have a similar niche breadth^20^. To evaluate the similarity of Zika and dengue, we built another niche model using the dengue occurrence database compiled by Messina *et al.*^21^. Comparing the two niche models reveals that the two niches are significantly different (Schoener’s *D* = 0.256; *p* < 0.02; Extended Data Fig. 6). While the two occupy a similar region of global climate space, Zika is more strictly tropical than dengue, occupying regions with higher diurnal temperature fluctuations and seasonality of precipitation (Fig. 2a). Moreover, our future projections for dengue (which strongly agree with previously published ones^22^) show an expansion out of the tropics that is not shared with Zika (Fig. 2, 3). These results call into question the applicability of dengue niche models used to project a significant future range for Zika in North America^6^.

**Figure 3.**
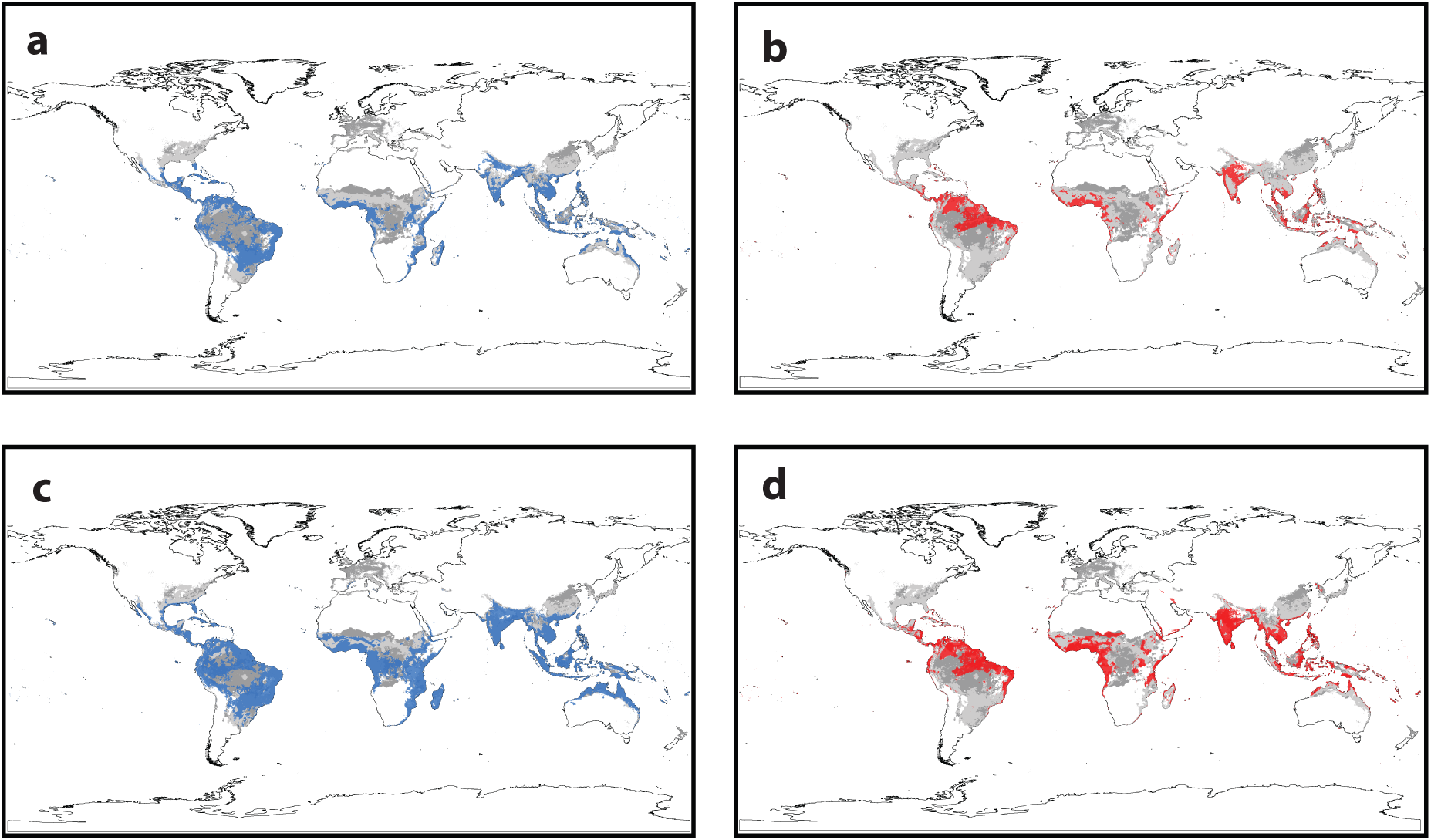
The estimated distribution of Zika (red) and dengue (blue) based on current (a, b) and 2050 climate projections (c, d), compared against the current (light grey) and future distribution of all three mosquito vectors (a-d).

Given the ecological nonequivalence of Zika and dengue, and the occurrence of Zika cases outside our predicted suitable range for the virus, the 2016 Zika outbreak may be in ephemeral rather than stable parts of the Zika transmission niche due to anomalous climatic conditions. Specifically, El Niño Southern Oscillation (ENSO) events drive outbreaks of dengue in the Americas and in Southeast Asia^23^, and we conjecture that the 2016 ENSO event may be responsible in large part for the severity of the ZIKV outbreak in North and Central America, a hypothesis also raised by Paz *et al.*^7^ in response to Bogoch *et al.*^6^. While wind-dispersed mosquitoes carrying infections can be responsible for the introduction of diseases to new regions^24^, reported cases in the United States have all been contracted sexually or while traveling abroad to regions with endemic outbreaks, further supporting the tropical constraint hypothesis. However, the rapid expansion during the current outbreak beyond the boundaries of the stable transmission niche indicates that regions outside our modeled range may support transmission during anomalous periods of climatic flux, but will not necessarily enable naturalization of the pathogen in the future. This highlights one of the most important limitations of this work, as ecological niche models relate occurrence to climate, while disease drivers may operate at the temporal scale of weather.

While the potential for rare, weather-driven outbreaks should not be overlooked, our models imply that it is premature to expect Zika naturalization as an eventuality in North America. Without more definitive information on the basic biology of Zika, however, the confidence with which niche models can forecast pandemics is limited. In particular, we draw attention to recent evidence suggesting Zika persistence may depend on wildlife reservoirs in addition to human hosts and mosquitoes. Primates have been suggested as the primary candidate clade, because the Zika flavivirus was first isolated in a rhesus macaque in the Zika Forest in Uganda. But as rhesus macaques do not occur on the African continent, and were captive there for inoculation experiments, the primate reservoir hypothesis remains unsupported. A 2015 case of an Australian presumed to have contracted Zika from a monkey bite while traveling in Indonesia, however, indicates that primates may transmit the virus directly^12^. Additionally, antibodies against Zika have been observed in several rodent and livestock species in Pakistan^25^, as well as several large mammal species, including orangutans, zebras, and elephants^26^. The potential for any North American wildlife species to play host to Zika is, at the present time, entirely unknown, and the infection of alternate hosts could potentially support new regions of stable transmission.

From the limited data in existence, we conclude that the global threat of a specifically vector-borne Zika pandemic, though devastating, may be limited to the tropics. However, sexual transmission of Zika infections may still facilitate a significant outbreak in the United States and other previously unsuitable regions, particular under evolutionary processes that select for the most directly transmissible strains of pathogens^27^. A case of sexual transmission in Texas has been suspected in the 2016 outbreak, and two previous reports of likely sexual transmission of ZIKV originate from 2011 and 2015^3,28^, though these seem to have been overlooked in most press coverage, which has presented the case of sexual transmission in Texas as a novel facet to the disease. Even if the Zika cases in the United States represent a rare spillover outside of the mosquito-borne viral niche, sexual transmission could create a new, unbounded niche in which the virus could spread. We draw attention to the potential parallels with simian and human immunodeficiency virus (SIV/HIV), for which a sexually transmitted pandemic has overshadowed the zoonotic origin of the disease^29^. With Zika’s asymptomatic presentation and the overall confusion surrounding its basic biology and transmission modes, we caution that its potential for a sexually-transmitted global pandemic cannot be overlooked in the coming months.

## Supplementary Information

Supplementary Information is linked to the online version of the paper at www.nature.com/nature.

## Acknowledgements

C.J.C. thanks Fausto Bustos for feedback on initial ideas presented in the paper, and Kevin Burgio for extensive methodological support, training and mentorship. We also thank two anonymous reviewers for their aid in strengthening the manuscript.

## Author Contributions

C.J.C. and E.R.D. collected the data, ran the models and wrote the first draft. All authors edited and approved the final text submitted for review.

## Author Information

Data presented in the paper are available in Table S1. Reprints and permissions information is available at www.nature.com/reprints. The authors declare no competing financial interests. Correspondence and requests for materials should be addressed to C.J.C.

## Methods

Occurrence data for Zika virus was compiled from the literature from studies dating as far back as the original discovery of the virus in Zika Forest, Uganda in 1947. Special attention was paid to correctly attributing cases of travelers to the true source of infection. Locality data was extracted from a set of clinical and survey papers, and georeferenced using a combination of Google Maps for hospitals and the Tulane University GEOLocate web platform for the remainder^1^, which allows for the attribution of an uncertainty radius to points only identified to a regional level. Sixty four points were used in the final models presented in our paper after limiting our results to only locations that could be estimated within 75 km (Extended Data: Table S1). To our knowledge, this spatially explicit database is the most inclusive dataset currently in the literature.

Occurrence data for the other species included in our study were compiled from the literature. For *Aedes africanus*, we used a dataset of 99 points downloaded from the Global Biodiversity Informatics Facility (www.gbif.org). GBIF’s coverage of *Aedes aegypti* and *Aedes albopictus* was however deemed to be lacking; occurrences for those species was taken from the previously published work of Kraemer *et al.*^2,3^ Messina *et al.*’s database was used for dengue^4^, as it has been previously published in *Scientific Data* and used with great success to generate a global distribution model.^5^ Both of these datasets were reduced down to point-only data (i.e., polygons of occurrence were excluded), leaving 5,216 points for dengue and 13,992 and 17,280 points for *Ae. aegypti* and *Ae. albopictus* respectively.

Due to the potentially transient nature of the New World distribution of Zika virus, our model uses presence and 1000 randomly selected pseudo-absence points from the Eurasian, African, and Australian regions where the virus is established. We used the WorldClim data set BIOCLIM at 2.5 arcminute resolution to provide all but one of our climate variables.^6^ The BIOCLIM features 19 variables (BIO1-BIO19) that summarize trends and extremes in temperature and precipitation at a global scale. Given the relevance of the normalized difference vegetation index (NDVI) in previous studies of dengue and as a predictor of vector mosquito distributions^7^, we downloaded monthly average NDVI layers for each month in 2014 from the NASA Earth Observations TERRA/MODIS data portal^8^, and averaged those twelve layers to provide a single mean NDVI layer. Species distribution models were executed using the BIOMOD2 package in R 3.1.1, which produces ensemble species distribution models using ten different methods: general linear models (GLM), general boosted models or boosted regression trees (GBM), general additive models (GAM), classification tree analysis (CTA), artificial neural networks (ANN), surface range envelope (SRE), flexible discriminant analysis (FDA), multiple adaptive regression splines (MARS), random forests (RF), and maximum entropy (MAXENT).^9^ Models were run individually for Zika (ZIKV), dengue (DENV), *Ae. aegypti, Ae. albopictus*, and *Ae. africanus*. For Zika, models trained on Old World environmental data were used to establish the potential distribution of the virus in the Americas under climatic conditions captured by WorldClim data, which represent an expected range of variability that do not incorporate anomalous events like 2015 El Niño Southern Oscillation.

To address colinearity in the environmental variable set, we produced a correlation matrix for our 20 variables, and identified each pair with a correlation coefficient > 0.8. For each species, we ran a single ensemble model with all ten methods and averaged the variable importance for our 20 predictors across the methods (See Table S2-S6). In each pair we identified the variable with the greater contribution, and we produced species-specific reduced variable sets used in the final published models by eliminating any covariates that universally performed poorer than their pairmate. Based in this criteria, we excluded the following variables for each species to reduce colinearity:

- ZIKV: BIO8, BIO9, BIO14, BIO18
- DENV: BIO3, BIO5, BIO12, BIO17
- *Ae. aegypti:* BIO6, BIO8, BIO12, BIO17
- *Ae. africanus:* BIO5, BIO6, BIO12, BIO17
- *Ae. albopictus:* BIO8, BIO9, BIO16, BIO17

The AUC of every model run with reduced variable sets is presented in Table S7. We found no significant correlation between NDVI and any individual BIOCLIM variable, so NDVI was included in every model of current distributions. We ran five iterations of each reduced variable set model and eliminated any prediction methods from the ensemble with an AUC of lower than 0.95, so that the final model had only included the best predicting models. This was found to only leave the RF method for DENV, so a cutoff of 0.9 was applied in that case, to keep the ensemble approach constant across datasets. The final models were run with the following methods with ten iterations using an 80/20 training-test split in the final presentation:

- ZIKV: GLM, GBM, GAM, CTA, FDA, MARS, RF
- DENV: GLM, GBM, GAM, FDA, MARS, RF, MAXENT
- *Ae. aegypti:* GLM, GBM, GAM, CTA, ANN, FDA, MARS, RF
- *Ae. africanus:* GLM, GBM, GAM, CTA, ANN, FDA, MARS, RF
- *Ae. albopictus:* GLM, GBM, GAM, CTA, FDA, MARS, MAXENT, RF

The importance of variables of the reduced model set for each are presented in Table S8-S12.

To project the distribution of the species under climate change, we reran each model with the previously chosen method and variable sets but excluding NDVI, for which we did not feel we could appropriately simulate future values. BioClim forecasts were taken from WorldClim using the Hadley Centre Global Environmental Model v. 2 Earth System climate forecast (HadGEM2-ES) predictions for representative climate pathway 8.5 (RCP85), which, within that model, represents a worst case scenario for carbon emissions and climate warming.^10^ All five species’ models were retrained on current climate data and projected onto forecasts for the year 2050, the results of which are shown in Figure 3. Finally, to compare the niche of dengue and Zika, we used the R package ecospat, which uses principal component analysis to define the position of species’ ecological niche relative to background environmental variation^11,12^. The ecospat analysis was run using the full 64 point database and the full extent of global environmental data, because, while the niche of Zika in the Americas is uncertain, dengue is well established, and measuring its niche required a full background sample. We excluded BIO5 and BIO12 from our analysis as they were included in neither of the final models for the diseases; niche similarity tests were run 100 times with 100 iterations each. The results of that analysis are presented in Figure S1, which shows both the one-directional similarity test and the bidirectional equivalence test.

Finally, to assess the transferability of our Zika model across environmental space, we conducted a geographic cross validation (GCV) between African and Asian datasets. While under normal circumstances, a model would be trained on New World data and projected onto the Old World, the lack of data prior to the current outbreak makes such a direct comparison infeasible. However, given the evidence for separate Asian and African strains, a cross-validation between the two was supported, and models trained on those two continents were projected globally to test the performance of the model across geographic regions (Extended Data Fig. 7), and evaluate how sensitive our projections in the Americas are to the environmental covariates sampled. The clustering of points in western India narrows the environmental range sampled by presences, significantly limiting the transferability of the Asian sub-model. In contrast, the African sub-model performs well in new regions, and corresponds well to the global model.

## Extended Data

**Extended Data Figure 1** | Final ensemble model for Zika virus

**Extended Data Figure 2** | Final ensemble model for dengue fever

**Extended Data Figure 3** | Final ensemble model for *Aedes aegypti*

**Extended Data Figure 4** | Final ensemble model for *Aedes africanus*

**Extended Data Figure 5** | Final ensemble model for *Aedes albopictus*

**Extended Data Figure 6** | Niche overlap between ZIKV and DEV

**Extended Data Figure 7** | Geographic cross validation

